# Deep learning inference of miRNA expression from bulk and single-cell mRNA expression

**DOI:** 10.1101/2025.05.03.652014

**Authors:** Rony Chowdhury Ripan, Tasbiraha Athaya, Xiaoman Li, Haiyan Hu

## Abstract

Understanding the activity of miRNA in individual cells presents a challenge due to the limitations of single-cell technologies in capturing miRNAs. To tackle this obstacle, we introduce two deep learning models: Cross-Modality (CM) and Single-Modality (SM). These models utilize encoder-decoder architectures to predict miRNA expression at the bulk and single-cell levels from mRNA data. We compared CM and SM with a state-of-the-art approach, miRSCAPE, using both bulk and single-cell datasets. We found that both CM and SM outperformed miRSCAPE in terms of accuracy. We also observed that integrating miRNA target information led to a significant enhancement in performance compared to using all genes. These models offer valuable tools for predicting miRNA expression from single-cell mRNA data.

## Introduction

MicroRNAs (miRNAs) are short, approximately 22 nucleotides in length, non-coding RNAs [1]. They commonly bind to the 3' untranslated regions of target mRNAs and control their expression through mRNA degradation and translation inhibition [2]. miRNAs play pivotal roles in dictating cellular destiny by governing crucial processes like development, maturation, differentiation, apoptosis, etc. [3]. Thus, miRNAs are intricately linked to various diseases, including cancer, periodontal disease, cardiovascular diseases, diabetes, and so on [3–6].

Single-cell technologies, particularly single cell RNA sequencing (scRNA-seq), have revolutionized biology by enabling the examination of molecular characteristics at the individual cell level, revealing heterogeneity and identifying novel cell types in diverse contexts [7]. However, the generation of miRNA profiles at single-cell resolution remains limited, primarily due to challenges in applying current sequencing protocols to miRNAs [8,9]. The scarcity of miRNA profiles at single-cell resolution hampers our understanding of miRNA functionality and dynamics at the individual cell level, as well as our exploration of miRNA involvement in disease development [2]. Therefore, it is crucial to predict miRNA expression at single-cell resolution from mRNA data.

There are only a few studies that infer miRNA expression from mRNA data. Chang et al. [10] proposed a seven-step computational approach leveraging motifs as potential miRNA target sites to predict miRNA expressions. Similarly, miREACT [11] employed motif enrichment analysis to infer miRNA activity from single-cell mRNA expression. Both methods were labor-intensive and dependent on motif analysis. A study conducted by Setty et al. [12] integrated mRNA expression, miRNA expression, copy number, and regulatory sequence data, utilizing a lasso regression model to predict miRNA expression. Additionally, Olgun et al. introduced miRSCAPE [2], which utilized an XGBoost classifier to predict miRNA expression from scRNA-seq data and showed superior performance to other methods. Despite the learning capabilities of these models, the accuracy of inference could potentially be enhanced by leveraging deep learning methods, given their superior feature learning and representation capabilities [1,13,14].

Recently, there has been a notable increase in the development of multimodal deep learning (MDL) techniques, facilitating single-cell multi-omics data integration and cross-modality prediction [15]. MDL methods can unveil intricate patterns and offer additional insight into the molecular characteristics of individual cells [16]. Unlike traditional computational methods for cross-modality prediction (such as matrix factorization [17,18] and correlation-based approaches [19,20]), which rely on manually extracted features for each data type, MDL can autonomously learn hierarchical representations for each modality by extracting meaningful features using a multilayer neural network (NN) models. In addition, MDL can handle high-dimensional data by mapping features from different modalities into a unified, smaller subspace [21]. For instance, BABEL [22] integrates scRNA-seq and single-cell ATAC-seq (scATAC-seq) data utilizing encoder-decoder NNs and predicts single-cell expression data. In addition, BABEL also facilitates cross-modality translation of scRNA-seq to scATAC-seq and scATAC-seq to scRNA-seq data.

In this study, we introduce two deep-learning models for inferring miRNA expression from both bulk and single-cell mRNA expression data. The Cross-Modality (CM) model, comprising two encoders and two decoders, predicts miRNA and mRNA expression using MDL techniques. The second model, an autoencoder referred to as SM, solely utilizes mRNA expression to infer miRNA expression. Evaluations on large cohorts from The Cancer Genome Atlas (TCGA) [23] program demonstrate the CM and SM models’ superior accuracy compared with a state-of-the-art machine learning approach, miRSCAPE [2]. Furthermore, leveraging bulk data for model training, both CM and SM exhibit better performance on single-cell datasets than miRSCAPE.

## Results

### CM and SM have a better performance on bulk data than miRSCAPE

We trained the CM model with 80% randomly selected paired miRNA-mRNA bulk data from the ten cancer types together and then assessed the model performance on the remaining 20% of bulk data from each of the ten cancer types separately (Table 1). We measured the correlation of the predicted miRNA expression and the actual miRNA expression across samples for each of the ten cancer types. Overall, CM achieved an average Spearman’s correlation of 0.58 and a Pearson’s correlation of 0.60 across the ten cancer types.

**Table 1.**
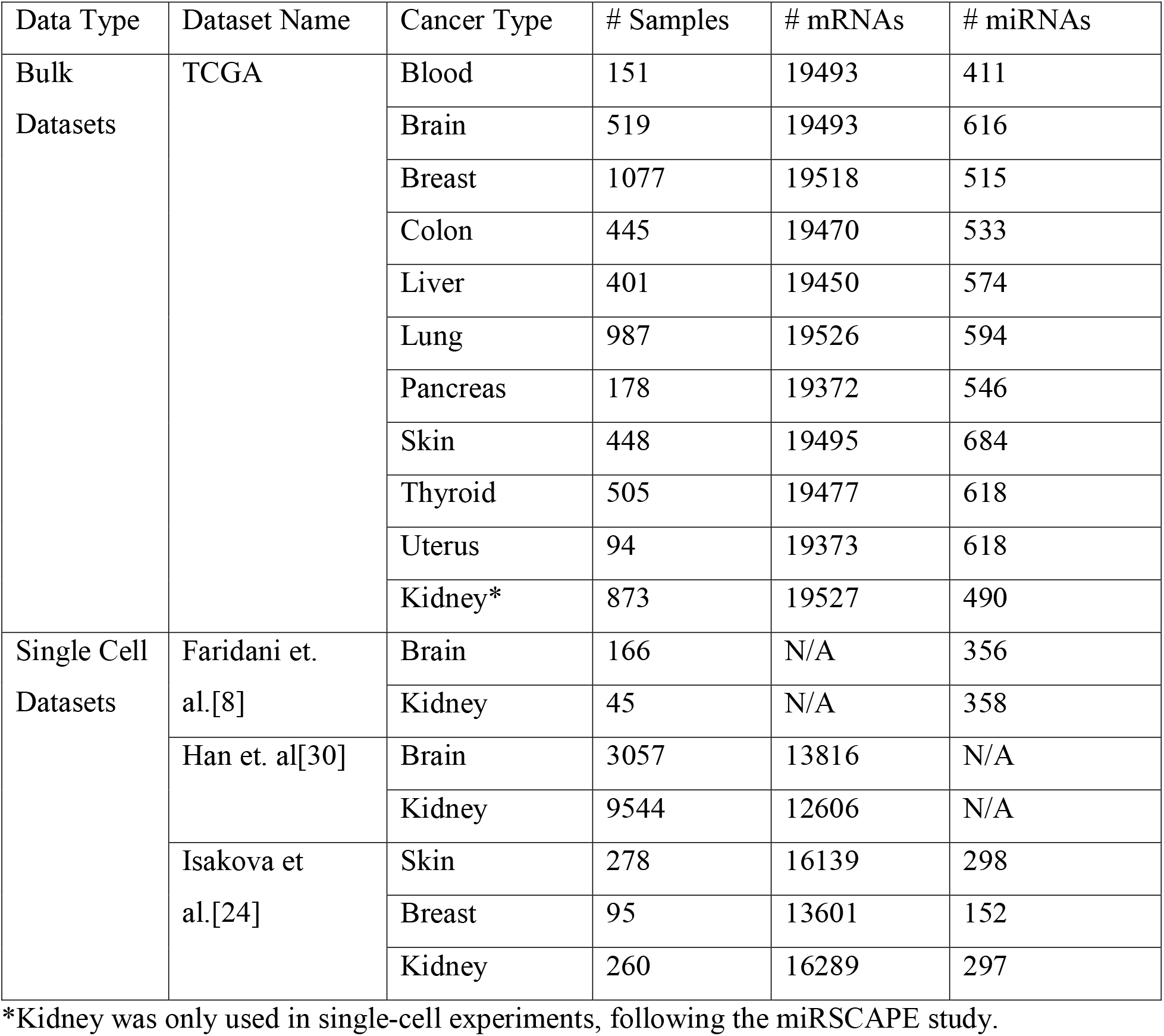
The number of samples, miRNAs and mRNAs considered.

| Data Type | Dataset Name | Cancer Type | # Samples | # mRNAs | # miRNAs |
| --- | --- | --- | --- | --- | --- |
| Bulk Datasets | TCGA | Blood | 151 | 19493 | 411 |
|  |  | Brain | 519 | 19493 | 616 |
|  |  | Breast | 1077 | 19518 | 515 |
|  |  | Colon | 445 | 19470 | 533 |
|  |  | Liver | 401 | 19450 | 574 |
|  |  | Lung | 987 | 19526 | 594 |
|  |  | Pancreas | 178 | 19372 | 546 |
|  |  | Skin | 448 | 19495 | 684 |
|  |  | Thyroid | 505 | 19477 | 618 |
|  |  | Uterus | 94 | 19373 | 618 |
|  |  | Kidney* | 873 | 19527 | 490 |
| Single Cell Datasets | Faridani et. al.[8] | Brain | 166 | N/A | 356 |
|  |  | Kidney | 45 | N/A | 358 |
|  | Han et. al[30] | Brain | 3057 | 13816 | N/A |
|  |  | Kidney | 9544 | 12606 | N/A |
|  | Isakova et al.[24] | Skin | 278 | 16139 | 298 |
|  |  | Breast | 95 | 13601 | 152 |
|  |  | Kidney | 260 | 16289 | 297 |

\*Kidney was only used in single-cell experiments, following the miRSCAPE study.

We studied how miRNA target information may improve the model performance. In one scenario, we considered all genes in the dataset. In the other scenario, we considered only genes that were targets of at least one of the miRNAs in the dataset. When considering all genes rather than only miRNA target genes, the CM model exhibited a reduced prediction accuracy across all cancer types (Material and Methods, Table 2). For instance, the average Spearman’s correlation decreased from 0.58 to 0.53, and the Pearson’s correlation dropped from 0.60 to 0.55, when incorporating all genes instead of solely focusing on miRNA target genes. This observation suggests that miRNA target genes provide valuable information for inferring miRNA expression. By focusing on target genes, we may have significantly reduced the unrelated genes and can thus model the miRNA expression better than considering all genes in the dataset.

**Table 2.** Comparison of the prediction accuracy of CM, SM, and miRSCAPE using all genes or only target genes across ten cancer types.

| Cancer Type | Spearman's correlation |  |  |  |  |  | Pearson's correlation |  |  |  |  |  |
| --- | --- | --- | --- | --- | --- | --- | --- | --- | --- | --- | --- | --- |
|  | CM |  | SM |  | miRSCAPE |  | CM |  | SM |  | miRSCAPE |  |
|  | all | targets | all | targets | all | targets | all | targets | all | targets | all | Targets |
| Blood | 0.50 | 0.58 | 0.48 | 0.56 | 0.40 | 0.40 | 0.52 | 0.62 | 0.50 | 0.60 | 0.43 | 0.44 |
| Brain | 0.63 | 0.70 | 0.63 | 0.69 | 0.53 | 0.58 | 0.65 | 0.72 | 0.66 | 0.71 | 0.57 | 0.60 |
| Breast | 0.47 | 0.52 | 0.47 | 0.52 | 0.45 | 0.47 | 0.50 | 0.54 | 0.50 | 0.54 | 0.48 | 0.48 |
| Colon | 0.50 | 0.56 | 0.51 | 0.56 | 0.43 | 0.45 | 0.51 | 0.56 | 0.51 | 0.56 | 0.44 | 0.46 |
| Liver | 0.56 | 0.60 | 0.55 | 0.62 | 0.48 | 0.51 | 0.58 | 0.62 | 0.57 | 0.64 | 0.51 | 0.53 |
| Lung | 0.53 | 0.57 | 0.52 | 0.57 | 0.49 | 0.51 | 0.54 | 0.57 | 0.53 | 0.58 | 0.50 | 0.51 |
| Pancreas | 0.46 | 0.50 | 0.44 | 0.51 | 0.37 | 0.40 | 0.48 | 0.53 | 0.47 | 0.53 | 0.39 | 0.40 |
| Skin | 0.57 | 0.60 | 0.56 | 0.59 | 0.50 | 0.48 | 0.58 | 0.62 | 0.58 | 0.61 | 0.52 | 0.50 |
| Thyroid | 0.48 | 0.56 | 0.50 | 0.57 | 0.43 | 0.46 | 0.5 | 0.57 | 0.52 | 0.58 | 0.44 | 0.46 |
| Uterus | 0.63 | 0.65 | 0.63 | 0.68 | 0.57 | 0.57 | 0.65 | 0.68 | 0.65 | 0.69 | 0.59 | 0.59 |
| Average | 0.53 | 0.58 | 0.53 | 0.59 | 0.47 | 0.48 | 0.55 | 0.60 | 0.55 | 0.60 | 0.49 | 0.50 |

We also conducted a comparison among the CM, SM, and miRSCAPE models using only miRNA target information. Across all ten cancer types, the CM and SM models consistently outperformed miRSCAPE (Figure 1, Table 2, Figure S1). Specifically, the CM model achieved an average Spearman’s correlation of 0.58, while the SM model attained an average of 0.59. In contrast, miRSCAPE attained an average correlation of 0.48 across the ten cancer types. Moreover, in terms of Pearson’s correlation, CM and SM both achieved an average correlation of 0.60, whereas miRSCAPE attained correlations of 0.5, across the same cancer types.

**Figure 1.**
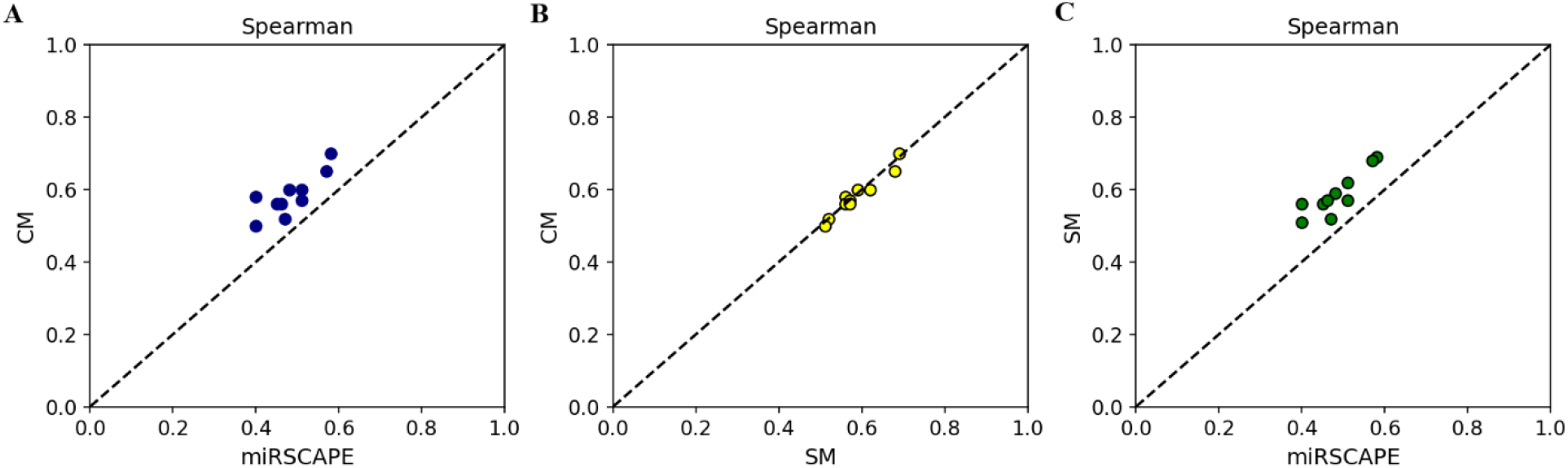
Scatter plot of the Spearman’s correlation between different models across ten cancer types incorporating only miRNA-mRNA target information. **A**. CM vs miRSCAPE; **B**. CM vs SM; and **C**. SM vs miRSCAPE.

We also did a five-fold cross-validation to compare the three models. Both the CM and SM models demonstrated strong performance, with an average Spearman’s correlation of 0.59 and Pearson’s correlation of 0.61 (Table S1, S2). In comparison, miRSCAPE achieved an average Spearman’s and Pearson’s correlation of 0.49 and 0.50, respectively (Table S3). It is worth noting that, CM has slightly better Spearman’s and Pearson’s correlation in each fold even though over five folds they have the same Spearman’s and Pearson’s correlation. In addition to the 80% training and 20% testing in the 5-fold CV, we also randomly divided our dataset into three parts, allocating 70% for training and randomly dividing the remaining data to create two test sets of equal size. In test set one, the CM model achieved an average Spearman's correlation of 0.58 and a Pearson's correlation of 0.60 across ten cancer types (Table S4). In test set two, the CM model achieved an average Spearman's correlation of 0.59 and a Pearson's correlation of 0.60 across ten cancer types (Table S4). SM performed similarly, while miRSCAPE showed smaller correlations. This comparison indicates the advantage of deep learning over classical machine learning approaches.

We also examined the performance of CM, SM, and miRSCAPE models prior to the inclusion of only miRNA target information (Table 2). Across all cancer types, the CM and SM models exhibited superiority over miRSCAPE with higher Spearman’s and Pearson’s correlations. Specifically, CM and SM achieved an average Spearman’s correlation of 0.53 while miRSCAPE trailed behind at 0.47. Similarly, regarding Pearson’s correlation, CM and SM both attained an average of 0.55, surpassing miRSCAPE 0.49. As observed for CM, the accuracy of both SM and miRSCAPE improved when we narrowed the input genes to only miRNA targets, the union of the target genes of all miRNAs considered in the dataset. For example, SM initially achieved Spearman’s and Pearson’s correlations of 0.53 and 0.55, respectively, which increased to 0.59 and 0.6 after considering only miRNA target data. Similarly, miRSCAPE had the Spearman’s and Pearson’s correlations of 0.47 and 0.49, respectively, with all genes in the dataset, which progressed to 0.48 and 0.5 after focusing on only miRNA targets. This underscores the potential for enhancing miRNA expression prediction accuracy through the inclusion of their target genes.

Subsequently, we evaluated the number of miRNAs with the Spearman’s correlation greater than 0.80 for each model across different cancer types (Figure 2). Overall, it is evident that CM and SM outperform the miRSCAPE model. For instance, for colon cancer, miRSCAPE had no miRNAs with the Spearman’s correlation greater than 0.80, whereas CM and SM had 4 and 1 miRNAs, respectively. For blood, brain, colon, skin, and thyroid, CM had more miRNAs with the Spearman’s correlation greater than 0.80 than SM. For liver, pancreas, and uterus, SM had more miRNAs than CM. This highlighted CM and SM’s superior prediction capability to miRSCAPE.

**Figure 2.**
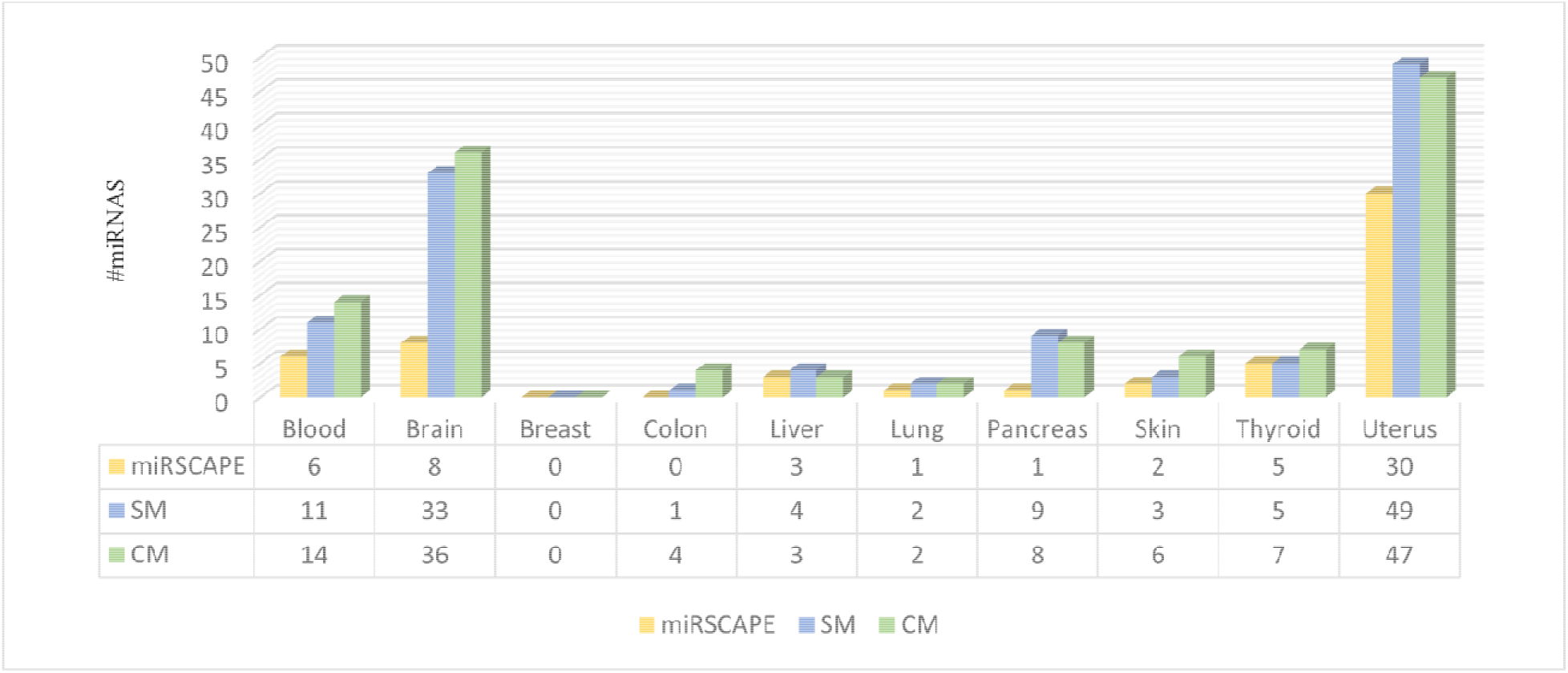
The #miRNAs with the Spearman’s correlation greater than 0.80 by the three models.

### CM and SM have a better performance on single cell data than miRSCAPE

After assessing the performance of CM and SM on bulk data, we evaluated their effectiveness in predicting miRNA expression on single-cell data. We utilized the dataset from Faridani et al. [8], which provide sc-miRNA expression profiles for brain (166 cells) and kidney (45 cells).

First, we trained CM, SM and miRSCAPE models using bulk brain and kidney cancer data from TCGA separately. We opted for such cancer-specific models trained on individual cancer type because single-cell data offers high resolution at the single-cell level, enabling detailed insights into cell-specific processes. Subsequently, we generated 50 pseudobulk samples for both the Faridani et al. and Han et al. data. A pseudobulk sample in a cell type was generated by randomly sampling 80% of the cells of this cell type with replacement and average the gene expression level of a gene in these 80% of cells as the expression level of this gene for every gene. The models were applied to the 50 generated pseudobulk mRNA samples to predict 50 miRNA samples. We then compared the predicted miRNA samples with the 50 generated pseudobulk miRNA samples by measuring the Spearman’s correlation of the log fold change (FC) of the predicted and observed expression of miRNAs. We found that SM exhibited a high Spearman’s correlation, 0.83, of the predicted log FC of miRNA expression and the observed log FC of miRNA expression between brain and kidney cells (Figure 3A), while CM and miRSCAPE achieved a Spearman’s correlation of 0.77 and 0.67, respectively. Notably, some miRNAs showed significant deviations between the predicted log FC and observed log FC (Figure 3A), which may be attributed to their low expression levels. In fact, approximately 98% of miRNAs with large deviations (absolute value of log FC difference >1) were relatively low-expressed miRNAs. In contrast, 81% of miRNAs with a high correlation between predicted and observed expression had relatively high expression. We also observed that certain miRNAs had near-perfect correlation, especially when the models were trained by cancer-specific training data (Figure 3), suggesting that cell-type-specific training data improved the model performance, and some miRNAs may have differential cell-type-specific expression.

**Figure 3.**
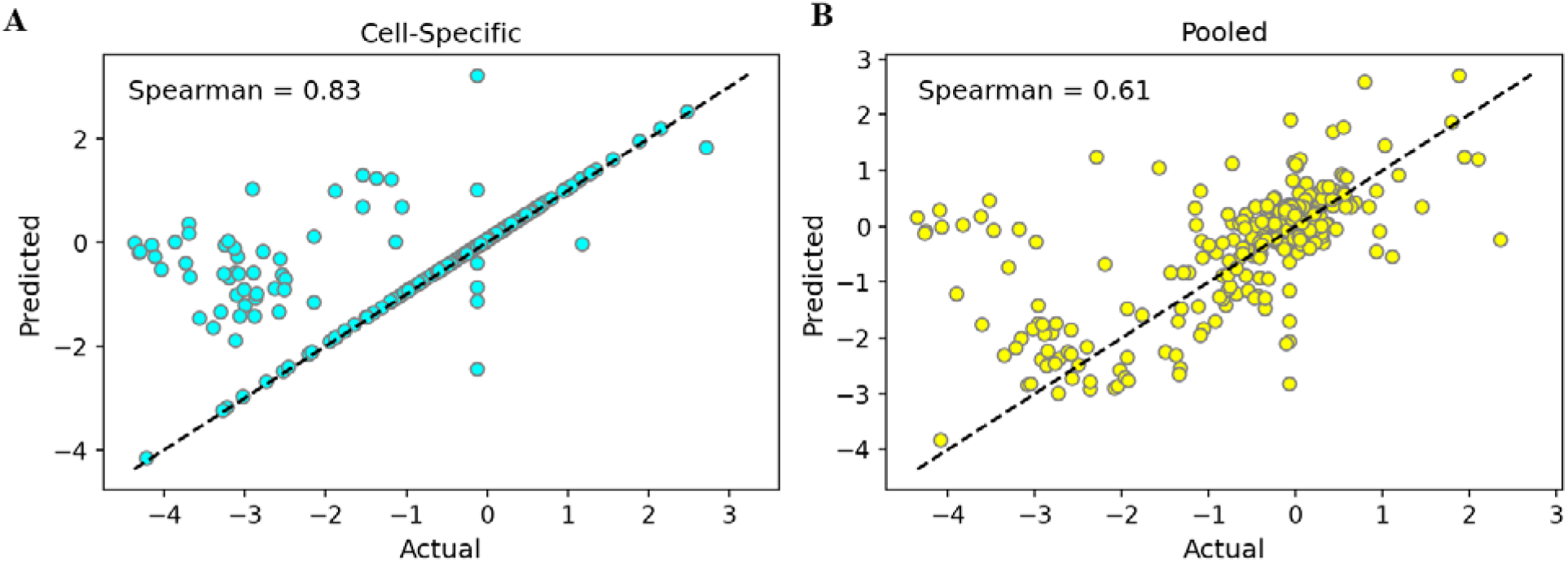
Evaluation on Faridani et al. data. Scatterplots representing comparison between observed miRNA’s log FC (X-axis) and predicted miRNA’s log FC (Y-axis). **A**. cell-specific model and **B**. a single pooled model.

Subsequently, we trained the models on pooled brain and kidney bulk data to predict miRNA expression for both kidney and brain single-cell data (Figure 3B). Once again, we compared the log FC between kidney and brain based on the predicted and observed miRNA expression from Faridani et al. CM achieved a high Spearman’s correlation of 0.61 (Figure 3B), while both SM and miRSCAPE obtained a correlation of 0.53. Similar to the above analysis on single-cell data from brain and kidney cells, we found that about 67% of miRNAs with large deviations between predicted and actual log FC were low-expressed, and 75% of the miRNAs with small deviations between predicted and observed log FC were more highly expressed.

Additionally, we utilized the dataset from Isakova et al. [24], which concurrently profiled mRNA-miRNA pairs within the same cell using Smart-seq-total in skin (278 cells), breast (95 cells), and kidney (260 cells). For each cell type separately, we trained our CM and SM models with cancer-specific TCGA bulk data and evaluated their performance on pseudobulk samples generated from single-cell mRNA (sc-mRNA) expression data to predict cell-type-specific miRNA expression. The correlation of log FC between the predicted and observed cell-type-specific miRNAs was assessed for each pair of cell types. When comparing “skin vs breast", “skin vs kidney", and “breast vs kidney", CM, SM, and miRSCAPE all exhibited a perfect Spearman’s correlation of 1.0. Similar to the observations in Figure 3, the perfect correlation was likely due to the cell-specific models trained.

We also applied the models to predict miRNA expression directly from sc-mRNA expression data instead of pseudobulk samples. In brief, for breast, kidney, and skin, we trained CM, SM, and miRSCAPE models with the TCGA bulk breast, kidney and skin cancer data, respectively, and evaluated their performance on sc-mRNA data from Isakova et al. to predict sc-miRNA expression. We then calculated the correlation of the log FC of the predicted and observed miRNA expression between the three pairs of cell types. Overall, CM and SM outperformed the miRSCAPE model for each pair of cell types (Table 3). In addition, CM performed better than the SM model in the “skin vs breast” and “skin vs kidney” comparisons. However, in the “breast vs kidney” comparison, SM (0.81) performed better than the CM (0.77) model.

**Table 3.** Comparison of the Spearman’s correlation of CM, SM, and miRSCAPE across different pairs of cell types at single-cell resolution.

|  | CM | SM | miRSCAPE |
| --- | --- | --- | --- |
| “breast vs kidney” | 0.77 | 0.81 | 0.68 |
| “skin vs breast” | 0.89 | 0.75 | 0.75 |
| “skin vs kidney” | 0.92 | 0.74 | 0.72 |

## Discussion

We introduced two deep learning models, CM and SM, to predict miRNA expression from mRNA expression. While the SM model focuses solely on inferring miRNA expression from mRNA expression, the CM model offers the added capability of cross-modality prediction, enabling the prediction of miRNA expression from mRNA expression and vice versa. We evaluated these models against miRSCAPE on both bulk and single-cell datasets. Overall, CM and SM surpassed miRSCAPE in performance on both data types. Note that when we trained and tested miRSCAPE, we used its default hyperparameters (Table S5), which may affect its performance.

Our study emphasizes the advantage of integrating miRNA target information, which has been shown to further boost the prediction accuracy compared with using all genes. This approach capitalizes on the specific regulatory roles of miRNAs, which are known to post-transcriptionally regulate gene expression by binding to target mRNAs, thereby influencing various cellular processes and disease states [25]. Focusing on miRNA target interactions helps reduce noise and irrelevant signals that may be present in broader gene-level data. By training on these specific interactions, models learn to identify and prioritize features directly involved in gene regulation, resulting in more effective learning and prediction of miRNA expression. Overall, the targeted feature selection not only reduces the dimensionality of the data, but also focuses on biologically relevant interactions, thereby improving the interpretability and accuracy of the models [26].

Although CM effectively leverages paired mRNA-miRNA expressions to infer miRNA expression from mRNA expression data, our models may encounter challenges in accurately predicting miRNA expression when confronted with significant data sparsity, particularly evident in single-cell datasets [27]. We observed that non-zero expression influences the prediction accuracy in our study. The developed CM and SM models cannot be directly applied to single-cell mRNA data to predict single-cell miRNA expression within individual cell types, because of the large variation of gene expression across cells within the same cell type. We thus applied the models on pseudobulk data from single-cell data instead of the original single-cell data and considered log FC of miRNA expression between a pair of cell types instead of within an individual cell type. In the future, we may enhance the robustness of CM and SM towards the sparsity inherent in single-cell datasets by employing deep count autoencoder [27] based models and other techniques to address this issue. Moreover, our approach to normalize library sizes before applying negative binomial (NB) distributions may not fully account for the increased variance in smaller libraries and introduce bias in data. Future work could explore utilizing sample-specific library sizes as a parameter for the NB distributions to improve prediction accuracy. In addition, although we tested CM and SM models on ten types of cancer and two single-cell datasets, it is necessary to further train and test the corresponding CM and SM models on other cancer and cell types, to experimentally validate the predictions, and to include more miRNAs in the study in order to understand the generality and portability of the models and advance their applications. Finally, we may improve the CM model by integrating mRNA-miRNA target information using a Graph Neural Network, which may allow for more precise modeling of mRNA-miRNA target relationships.

## Material and Methods

### Bulk and Single-cell Data

We assessed the performance of CM and SM using both bulk and single-cell datasets and compared them with miRSCAPE [2]. For a fair evaluation, we gathered RNA-seq data of paired miRNA-mRNA samples for the same ten cancer types from TCGA previously used by miRSCAPE. Paired here means that for each sample, both miRNAs and mRNAs were sequenced from the same type of tissues, not necessary from exactly the same tissue sample. Each of these ten cancer types contained 1881 miRNAs and 19938 mRNAs. Same as miRSCAPE [2], we removed mRNAs with zero expression in all samples and miRNAs expressed in fewer than half of the samples within each cancer type, which attempted to ensure the inclusion of relevant miRNAs and to have enough data for model training [28,29]. In this way, 31.62%, 40.37%, 32.25%, 34.48%, 37.49%, 36.51%, 37.76%, 42.75%, 39.14%, and 42.5% of miRNAs were kept in Blood, Brain, Breast, Colon, Liver, Lung, Pancreas, Skin, Thyroid, and Uterus cancer, respectively (Table 1).

For single-cell data analysis, we used two datasets from Faridani et al. [8] and Isakova et al. [24]. The Faridani et al. dataset consists of sc-miRNA expression profiles for 585 miRNAs in brain (166 cells) and kidney (45 cells). Since this dataset lacks paired sc-mRNA expression profiles, we obtained brain and kidney sc-mRNA data from Han et al.[30]. In the brain sc-mRNA dataset, there were 3057 cells and 15477 mRNAs, while in the kidney sc-mRNA dataset, there were 9544 cells and 18790 mRNAs. Isakova et al. [24] concurrently profiled mRNA-miRNA pairs within the same cell using Smart-seq-total in skin (278 cells), breast (95 cells), and kidney (260 cells). They identified 1683 sc-miRNAs and 18135 sc-mRNAs across these cell types.

### CM Model

#### Architecture and Design

The CM model consists of four modular NNs: two encoders and two decoders (Figure 4A). The two encoders are trained on either mRNA or miRNA expression data to create two 16-dimensional joint embeddings. Each joint embedding is then decoded into both miRNA and mRNA expression by the same two decoders. Note that the decoder to decode mRNA embedding and miRNA embedding into the expression of one miRNA at a time for the more accurate predicted miRNA expression. The four NNs are trained simultaneously to minimize the sum of four types of losses: the loss of the predicted mRNA expression from the mRNA embedding, the loss of the predicted *i-*th miRNA expression from the mRNA embedding, the loss of the predicted mRNA expression from the miRNA embedding, and the loss of the predicted *i-*th miRNA expression from the miRNA embedding. We used the NB loss for mRNA expression and the root mean squared error (RMSE) loss for miRNA expression. This choice of such loss functions for mRNA and miRNA stems from the different pipelines to process mRNA and miRNA data [31]. We also tested other options, including the RMSE loss for both mRNA and miRNA expression, the NB loss for both mRNA and miRNA, etc. We found that the choice of the NB loss for mRNA expression and the RMSE loss for miRNA expression resulted in the best performance.

**Figure 4.**
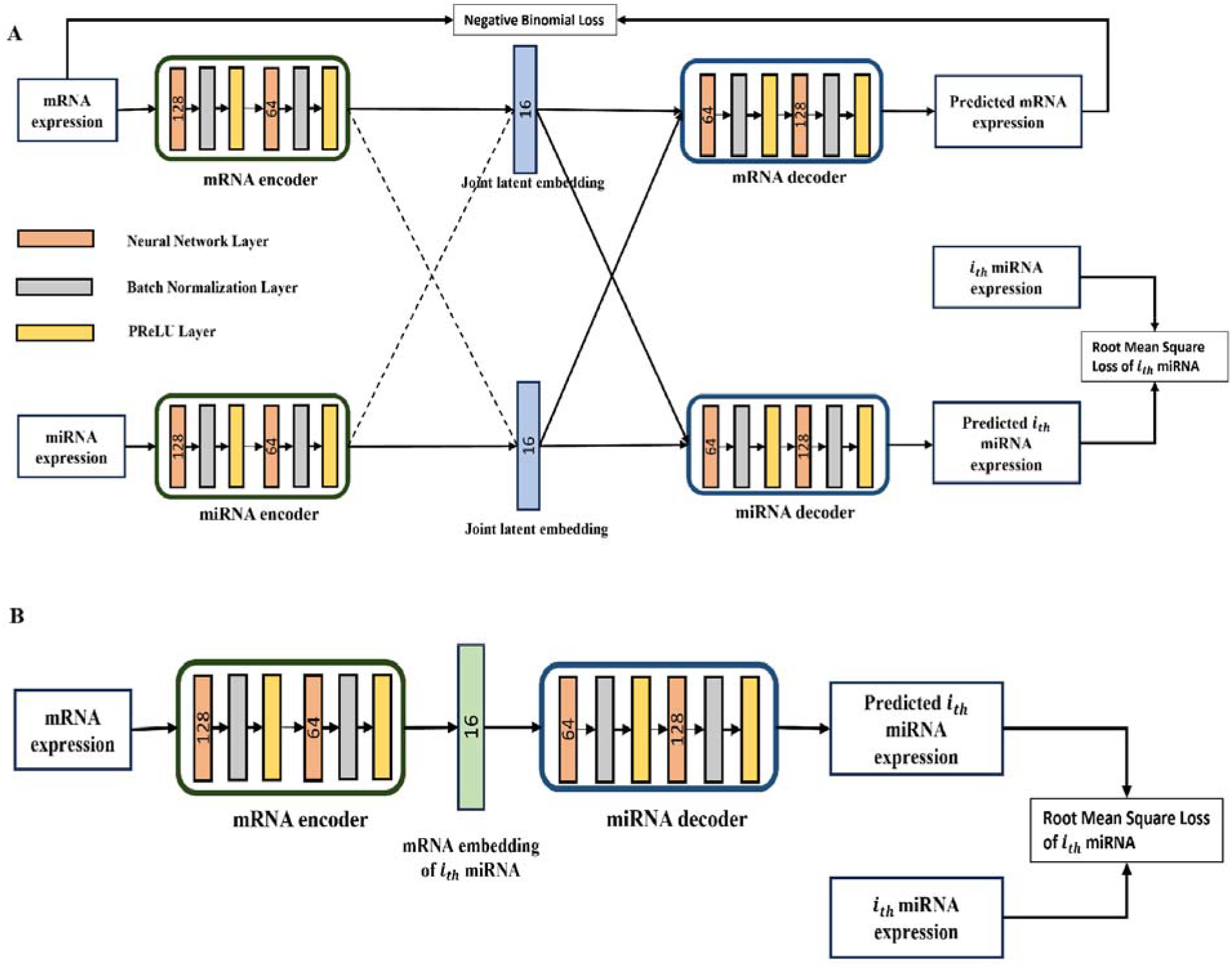
A graphical overview of the **A**. CM and **B**. SM models.

#### Implementation

The encoders and decoders consist of fully connected layers. Each of the two encoders has three layers, with 128, 64, and 16 neurons, respectively. Following each layer is a batch normalization layer and a parametric rectified linear unit (PReLU) layer. Similarly, each decoder also features three layers, with dimensions of 16, 64, and 128, respectively. The mRNA and miRNA decoders are succeeded by a batch normalization layer and a PReLU layer, to convert the 16-dimensional latent representation into 64 dimensions. Then, this 64-dimensional layer is projected into 128 dimensions, followed by a batch normalization layer and a PReLU activation function. For the miRNA decoder, the 128 dimensions are mapped to a single neuron to describe the expression of one miRNA. For the mRNA decoder, the 128 dimensions are mapped to two outputs, matching the size of the input mRNA expression vector. The output undergoes exponential and softplus activation functions to estimate the mean and dispersion of the expression. These parameters collectively characterize the probability of observing the expression of each gene according to a NB distribution. Coupled with a NB loss function, this enables the CM model to learn and estimate the actual, “de-noised” mRNA expression values (represented by the mean parameter), rather than the observed noisy values.

#### Loss Function

As mentioned above, the mRNA decoder outputs a pair of vectors, representing the mean nd dispersion () of the estimated expression, respectively, for each gene. These vectors are used to parameterize the probability of observing the measured mRNA expression () following a NB distribution, as illustrated below (where represents the Gamma function):

Now our goal is to identify the mean) and dispersion values that maximize the probability of the observed data. This is essentially the same as minimizing the negative logarithm of the likelihood, which we adopt as the loss function to evaluate predicted mRNA expressions below (*ϵ* can be interpreted as a pseudo-count, which is analogous to adding a small count to each observed value to avoid zero counts and stabilize the calculations):

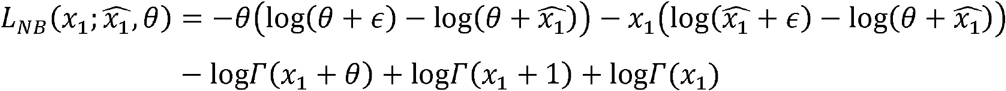

The RMSE loss is utilized to model the loss between the i_th_ predicted 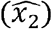 and observed (*x*_2_) miRNA expression data as in the following, where N is the number of samples in the dataset.

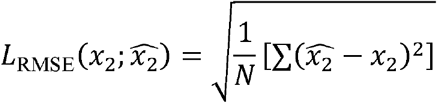

With these components in place, we have the following loss function L for our CM model:

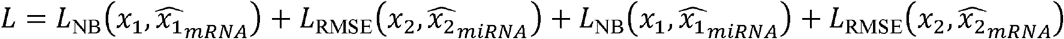

Here, *x*_1_ and *x*_2_ represent the observed mRNA and miRNA expressions, respectively; 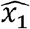 and 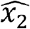 represent the predicted mRNA and miRNA expressions; and subscripts denote the source modality used to infer either mRNA or miRNA. For instance, 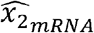 represents the predicted miRNA expression from mRNA input. The first two terms in the above loss function capture the CM’s ability to reconstruct the mRNA and miRNA expression through the intramodality inference. The last two terms capture the CM's performance in cross-modality prediction, meaning its capability to infer miRNA profiles from mRNA expression, and vice versa.

We employed the Adam optimizer during training. The hyperparameters were determined through an empirical trial-and-error process. Specifically, we evaluated four different learning rates (0.01, 0.001, 0.0001, and 1e-6), and four batch sizes (32, 64, 128, and 512). The combination of a learning rate of 0.01, and a batch size of 64 consistently yielded optimal performance for the CM model. To enhance training stability, we implemented gradient clipping [32]. This technique constrains the magnitude of gradients during backpropagation, preventing the occurrence of exploding gradients and ensuring controlled updates to model parameters [32].

### SM Model

#### Architecture and Design

Our SM model consists of two modular NNs: one encoder and one decoder (Figure 4B). The encoder is trained to project mRNA expression into a 16-dimensional latent representation for each miRNA. This latent representation works as a reduced feature dimension of mRNA profiles. The decoder then predicts the expression of this miRNA from the mRNA embedding.

#### Implementation

The encoder and decoder in SM consist of three fully connected layers. At the first layer, the encoder takes the input feature vector and projects it into 128 dimensions; in the second layer, the 128 dimension representations is transformed into 64 dimension ones; in the final layer, these 64 dimension representations are projected into 16-dimensional latent representations. All three layers are followed by a batch normalization layer and a PReLU nonlinear activation layer. The first two layers of our decoder fully invert the last two layers of the encoder. There is only one neuron at the last decoder layer to describe the expression of one miRNA.

#### Loss Function

For each miRNA, we train the SM model to predict its expression based on the global mRNA profile of the sample. For example, if we have 5 miRNAs, our SM model is trained five times with the same training mRNA data. At each training, Our SM model aims to predict expressions across samples for one miRNA. Then we calculate the RMSE loss between the predicted and observed miRNA expression and backpropagate the loss. Similar to CM, for each training, the Adam optimizer is utilized with a batch size set to 64 and a learning rate of 0.01.

#### Training and testing

Deep learning models require a substantial amount of data for effective training and accurate inference. Therefore, both the CM and SM models were trained using the combined data from the ten cancer types. Specifically, 80% of the data from each cancer type was randomly selected, combined, and used for training, while the remaining 20% was set aside for evaluation. The combined dataset comprised 19,257 mRNA entries and 356 miRNA entries. We also used these training and testing data to train and test miRSCAPE.

Given the pivotal role of miRNAs in regulating numerous human mRNAs and their involvement in various developmental and physiological processes [25], we hypothesized that incorporating miRNA target information could improve miRNA expression prediction accuracy. Consequently, we focused on mRNAs targeted by miRNAs to model miRNA expression, utilizing the conserved targets of miRNAs available in the TargetScanHuman 8.0 database [33]. This refinement resulted in a dataset consisting of 3,923 mRNAs and 163 miRNAs. The CM, SM, and miRSCAPE models based on miRNA targets were trained and tested with these filtered mRNAs and miRNAs.

For both CM and SM models, initially, mRNA data underwent size normalization so that the sum of expression of all genes in each sample is equal to the median of the sum of gene expression across all samples. Subsequently, the size-normalized expression was log-transformed and standardized to have a mean of zero and a variance of one. Additionally, expression values falling below the 0.5^th^ percentile of the data were replaced by the 0.5^th^ percentile value and any expression values exceeding the 99.5^th^ percentile will be replaced by 99.5^th^ percentile value.

For single-cell data, we utilized bulk data of individual cancer types from TCGA to train the model. We then tested the trained model with pseudobulk data generated from single-cell data of the corresponding cancer types. In brief, we generated pseudobulk data from single-cell data by bootstrapping [34]. For each single cell type, we generated 50 bootstrapped pseudobulk datasets by randomly sampling 80% of the cells with replacement and averaging their expression values as the corresponding gene expression values. We utilized this pseudobulk data as a test dataset for our CM and SM models on single-cell data. To mitigate technological differences between bulk and pseudobulk data, we ensured that the feature values were normalized within the range of 0 to 1 in each sample, both during model training and testing on the pseudobulk sc-mRNA data.

To assess the performance of our models on bulk data, we compute both Spearman’s and Pearson’s correlations between the predicted and observed expressions for each miRNA across all test samples. We compute only Spearman’s correlation on single-cell data because scRNA-seq data often exhibits higher noise and more outliers compared with bulk RNA-seq data [35]. The Pearson’s correlation, which is sensitive to noises and outliers, might not provide a robust measure of association in the presence of such noise and outliers [36]. To calculate the Spearman’s correlation on single-cell data, we estimate the FC of expression for each miRNA across a pair of cell types using the predicted and observed miRNA expression values and then calculate the Spearman’s correlation of two log FCs of miRNAs. In brief, for a miRNA, we calculated the first FC as the ratio of its observed expression in cell type one to its observed expression in cell type two. The observed expression of a miRNA in a cell type is the average expression of this miRNA in all cells in this cell type. We then calculate the second FC as the ratio of its predicted expression in cell type one to its predicted expression in cell type two. The predicted expression of a miRNA in a cell type is the predicted expression of a miRNA by the CM, SM, or miRSCAPE models with the input pseudobulk mRNA expression in this specific cell type. With the two log FC for each miRNA, we then calculate the Spearman’s correlation these log FC across miRNAs in a single-cell dataset to assess how well a model predicts the miRNA expression on this dataset. Note that in the above analysis, we may directly calculate the Spearman’s correlation of the observed and predicted expression of a miRNA across pseudobulk samples. However, the scmiR-seq data are too sparse and have large variations of miRNA expression. Such a Spearman’s correlation is basically around 0 for every miRNA from every model we have tried, including the miRSCAPE model. We thus follow the miRSCAPE study to measure the Spearman’s correlation of log FC.

## Data availability

All data and code are available at https://www.cs.ucf.edu/∼xiaoman/tools/miRExp/.

## Credit author statement

**Rony Chowdhury Ripan:** Formal analysis, Investigation, Methodology, Software, Validation, Visualization, Writing-original draft. **Tasbiraha Athaya:** Investigation, Validation, Visualization, Writing-review & editing. **Xiaoman Li:** Conceptualization, Funding acquisition, Project administration, Supervision, Validation, Writing-review & editing. **Haiyan Hu:** Conceptualization, Funding acquisition, Project administration, Supervision, Validation, Writing-review & editing.

## Acknowledgments

We want to thank the National Science Foundation for the financial support.

